# Nr2f1 controls mitochondrial dynamics in mouse adult-born hippocampal neurons

**DOI:** 10.1101/2022.03.25.485113

**Authors:** Bonzano Sara, Michelon Filippo, Crisci Isabella, Molineris Ivan, Neri Francesco, Oliviero Salvatore, Beckervordersandforth Ruth, Lie Dieter Chichung, Peretto Paolo, Bovetti Serena, Studer Michèle, De Marchis Silvia

**Author notes:** **Co-corresponding authors:** Sara Bonzano, Silvia De Marchis.

## Abstract

The nuclear receptor Nr2f1 plays a key role during cortical development by acting as a strong transcriptional regulator in embryonic and postnatal neural cells. In humans, its mutations cause the Bosch-Boonstra-Schaaf optic atrophy-intellectual syndrome (BBSOAS), a rare neurodevelopmental disorder characterized by multiple clinical features including optic nerve atrophy, intellectual disability, and autistic traits. In this study by genome-wide and *in silico* analyses we found that key factors for mitochondrial function and dynamics represent potential genomic targets under direct Nr2f1 transcriptional control in neurons. Although mitochondrial dysfunction is increasingly implicated in neurodevelopmental disorders, whether and how Nr2f1 can regulate mitochondria function in neural cells is still completely unknown. To address these questions, we have combined mouse genetics, neuroanatomical and imaging approaches. To genetically manipulate Nr2f1 function in newborn neurons we focused on the adult mouse hippocampal dentate gyrus, a site of persistent neurogenesis where Nr2f1 is highly expressed. We showed that conditional Nr2f1 loss-of-function within the hippocampal neurogenic niche lead to a reduced mitochondrial mass associated to mitochondrial fragmentation in newborn neurons, likely reflecting alteration in mitochondrial dynamics. Accordingly, we showed that the fusion factor Mfn2, which is as a putative Nr2f1 target, is downregulated following both Nr2f1 adult depletion and embryonic haploinsufficiency in mice. Overall, our study provides the first evidence of a crucial role of Nr2f1 in shaping mitochondria in neurons and opens a promising avenue for the identification of new mechanisms implicated in BBSOAS pathogenesis involving mitochondrial dysfunction.

## Introduction

The role of the key transcriptional regulator Nr2f1 (Nuclear receptor subfamily 2, group f, member 1), also known as COUP-TFI (Chicken Ovalbumin Upstream Promoter-Transcription Factor 1), has been intensively investigated during brain development, where it exerts pleiotropic functions including the control of neural stem cell competency, cortical area and cell-type specification, and axonal pathfinding (Bertacchi et al., 2019b). In humans, *NR2F1* has recently emerged as a disease gene: multiple *NR2F1* pathological variants cause the Bosch-Boonstra-Schaaf optic atrophy-intellectual syndrome (BBSOAS; OMIM: 615722), a rare monogenic autosomal-dominant disorder characterized by multiple clinical features including global developmental delay, mild-to-severe intellectual disability (ID), optic nerve atrophy, seizures and traits characteristic of the autism spectrum disorder (ASD) (Bosch et al., 2014; Chen et al., 2016; Kaiwar et al., 2017). Most variants in the NR2F1 gene of patients described so far are deletions and/or mutations predominantly located in the DNA-binding domain (DBD) and lead to haploinsufficiency or dominant-negative effects, thus compromising and/or completely abolishing NR2F1 transcriptional regulatory activity (Billiet et al., 2021; Bosch et al., 2016; Chen et al., 2016; Kaiwar et al., 2017; Rech et al., 2020).

Several neurological symptoms - such as optic atrophy, hypotonia, and seizure -described in BBSOAS patients are often associated to mitochondrial dysfunction in the nervous system (Frye, 2020; Lenaers et al., 2021; Valenti et al., 2014; Zsurka and Kunz, 2015). Nevertheless, no data are available on possible mitochondrial implications in the neurological symptoms of BBSOAS patients and whether and how NR2F1 can impact on mitochondrial dynamics and/or function in neural cells is still unknown. To address these questions, we exploited adult neurogenesis in the mouse hippocampal dentate gyrus (DG) to conditionally manipulate Nr2f1 in adult neural stem/progenitor cells (aNSPCs) (Bonzano et al., 2018). Indeed, in the DG subgranular zone (SGZ), aNSPCs ensure the continuous generation of new neurons whose integration into the preexisting circuitries is crucial for proper adaptive behaviors and cognitive functions (Aimone et al., 2014). Interestingly, DG immature neurons show higher Nr2f1 expression (Artegiani et al., 2017; Bonzano et al., 2018) and an overall increase in mitochondrial mass compared to aNSPCs (Beckervordersandforth, 2017; Steib et al., 2014). Moreover, Nr2f1 drives aNSPCs toward a neuronal lineage inhibiting an astroglial fate (Bonzano et al., 2018), and mitochondrial dysfunction impairs DG neurogenesis (Beckervordersandforth, 2017; Beckervordersandforth et al., 2017; Khacho et al., 2016; Steib et al., 2014). Thus, adult neurogenesis represents an excellent model to dissect the role of Nr2f1 on mitochondria.

In this study, we demonstrate that Nr2f1 controls mitochondrial mass and shape in adult-born DG neurons. By genome-wide and *in silico* analyses we provide evidence that this occurs through the regulation of nuclear genes encoding proteins essential for mitochondrial dynamics and function. Among them, we identified the mitochondrial membrane protein mitofusin-2 (Mfn2) that plays a central role in regulating mitochondrial fusion, as a putative target gene of Nr2f1 transcriptional activity. We found that Mfn2 is downregulated in the DG of mice heterozygous for Nr2f1, a validated BBSOAS mouse model (Bertacchi et al., 2019a; Chen et al., 2020; Jurkute et al., 2021), and we confirmed its downregulation in adult-born DG neurons conditionally depleted for Nr2f1, supporting a cell-intrinsic effect. Overall, our data provide new insights into the functional role of Nr2f1 by demonstrating its direct involvement in shaping mitochondria in adult-born DG neurons. This paves the way for further research on the mechanisms and role of mitochondrial dysfunction in the pathogenesis of BBSOAS, opening new possibilities for therapeutic intervention.

## Results and Discussion

### Nr2f1 nuclear genomic targets are enriched in key mitochondrial factors

To investigate whether the nuclear transcription factor Nr2f1 impinges on mitochondria by directly binding and regulating the expression of nuclear-encoded mitochondrial genes that code for more than 95% of the mitochondrial proteins (Calvo et al, 2016; Misgeld & Schwarz, 2017) we run a genome-wide analysis of Nr2f1 occupancy by Chromatin immunoprecipitation followed by deep sequencing (ChIP-seq) (**Figure 1a**) on the adult neocortex, a brain region that is highly enriched in Nr2f1 expressing neurons (**Figure S1a,b**) (Foglio et al., 2021). By peak calling analysis, we identified 2119 binding sites for Nr2f1 and revealed that these sites were enriched in both CpG islands and promoter regions (i.e., −3kb/+2kb from the Transcription Starting Site - TSS - of annotated genes) (**Figure 1b and Figure S1c**). Almost all Nr2f1-bound promoters were positive for H3K4me3, a histone mark highly enriched at active promoters (Liang et al., 2004), but not all H3K4me3+ promoters turned to be bound by Nr2f1 (**Figure 1a**), implying specificity. Notably, predicting TF binding by the Homer motif discovery tool confirmed that the observed genomic peaks are highly enriched in the putative Nr2f1 consensus sequence (**Figure 1c and Figure S1d**), indicating direct binding. Together, these data show that in neurons the transcription factor Nr2f1 binds to promoters predominantly in a chromatin permissive state, in line with a previous study reporting a high association of Nr2f1/2 with open chromatin characterized by high levels of p300 and H3K27ac in neural crest cells (Rada-Iglesias et al., 2012).

**Figure 1.**
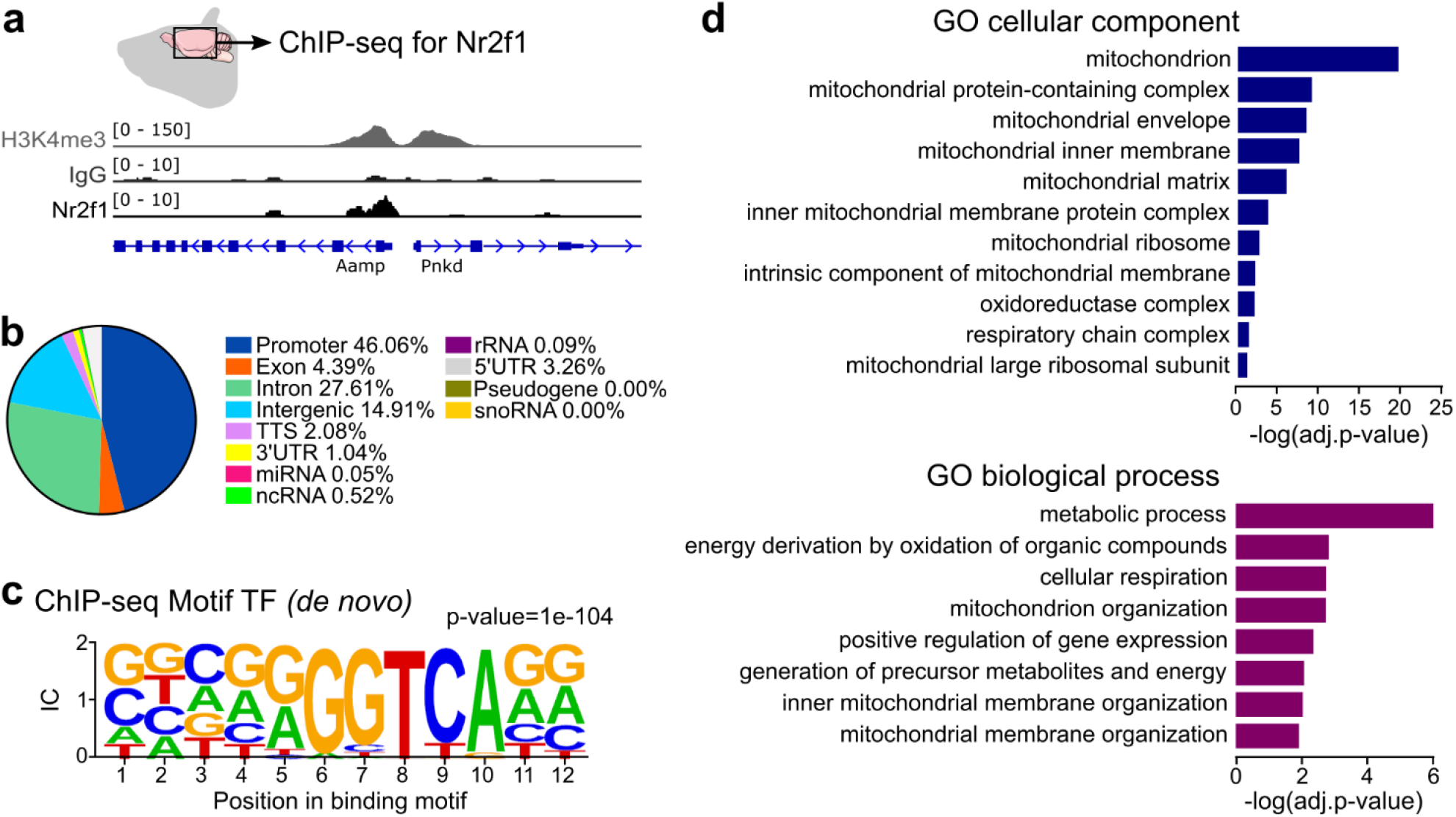
ChIP-seq analysis on adult mouse neocortex revealed direct binding of Nr2f1 on mitochondrial genes. **a.** Example of a genomic peak on a promoter showing Nr2f1 binding associated to the H3K4me3 (gene on the left) compared with a promoter region showing H3K4me3 without Nr2f1 binding (gene on the right). **b.** Pie chart illustrating the relative distribution (expressed as percentage) of the 2119 Nr2f1-bound sequences. **c.** De novo motif discovery of TF motif in Nr2f1 binding sites (forward) – see Figure S1e for matches to known motifs. **d.** Bar graph illustrating gene ontology (GO) term analysis wherein the 2119 genes bound by Nr2f1 were ranked for cellular components (top) and biological processes (bottom). The full lists of enriched categories are listed in Supplementary Table S1.

The set of genes identified as putative Nr2f1 genomic targets by ChIP-seq was then analyzed by Gene Ontology (GO) term annotation through the PANTHER classification system. Concerning the “cellular component” class, the analysis revealed a significant enrichment in genes involved in the control of mitochondrial structure (e.g. mitochondrial membranes and matrix organization) (**Figure 1d**). Notably, GO clustering of Nr2f1-bound genes according to the cellular localization of their products clearly showed a higher fold-enrichment and more significant over-representation for mitochondria (e.g., “mitochondrion”, fold-enrichment=2.04; FDR=8.20e-21) rather than non-mitochondrial subcellular location (e.g., “cytoskeleton”, fold-enrichment=1.27, FDR=0.0425; “Golgi apparatus”, fold-enrichment=1.41, FDR=0.0061). In line, analysis of the “biological process” class revealed an enrichment in genes involved in mitochondrial function, including the organization of mitochondrial membranes as well as mitochondria-related metabolism and cellular respiration (**Figure 1d**). Interestingly, a previous study reported altered expression of nuclear genes encoding mitochondrial proteins (e.g., *SLC25A1, ECHS1, MRPL45, TIMM23*) in the *Nr2f1-null* embryonic neocortex (Montemayor et al., 2010). Moreover, by interrogating a recently published RNA-sequencing dataset (Chen et al., 2020), we found altered expression of multiple genes coding for mitochondrial proteins (e.g., *OPA1, GDAP1, ACO2, ATP5G2*) in the hippocampi of adult *Nr2f1-heterozygous* mice. Taken together, these data suggest that Nr2f1 may exert a direct role on the transcriptional control of nuclear-encoded mitochondrial genes in neurons.

### Nr2f1 manipulations in adult-born hippocampal neurons lead to altered mitochondrial mass and morphology

To study the role of Nr2f1 on mitochondria, we focused on the neurogenic niche of the adult hippocampus by exploiting an *in vivo* conditional loss-of-function (LOF) approach targeting the aNSPC lineage *via* tamoxifen (TAM) administration in triple transgenic mice carrying the inducible *Cre-recombinase* (*CreERT2*) under the Glast promoter, both Nr2f1 alleles flanked by loxP sites and the reporter gene encoding the yellow fluorescent protein (YFP) for lineage tracing (herein named *Nr2f1-icKO*; **Figure 2a**) (Armentano et al., 2007; Bonzano et al., 2018). Animals carrying both *Cre* and YFP alleles but wildtype for *Nr2f1* were used as controls (*Ctrl*). To label mitochondria in adult-born hippocampal neurons, a retroviral vector encoding the mitochondria-targeted fluorescent protein DsRed (RV-mitoDsRed) was stereotaxically injected into the DG two weeks after induction by TAM. Mice were then analyzed at 17 days post viral-injection (**Figure 2a’**), allowing recombined and transduced newborn cells to develop into doublecortin-expressing neurons (i.e., DCX+mitoDsRed+YFP+ cells). Notably, while in *Ctrl* mice YFP+ cells were virtually all Nr2f1-positive, in *Nr2f1-icKO* mice the population of YFP+ cells - including the specific subset transduced by the mitoDsRed virus - were substantially devoid of Nr2f1 staining (**Figure 2b,c**), thus validating *Nr2f1* inactivation in the aNSPC neuronal lineage of *Nr2f1-icKO* mice. To perform the analyses, we selected DCX+mitoDsRed+YFP+ newborn neurons from *Nr2f1-icKO* and *Ctrl* mice showing comparable patterns of the dendritic trees throughout the DG layers (i.e., the primary dendrite branching in the outer part of the granule cell layer and the dendritic arborization spanning the molecular cell layer; **Figure S2a and Figure 3a,b**). Morphometric analyses on those cells revealed no changes in the size of perikarya (**Figure S2a’**) but a decrease in the total dendritic length and in the number of dendritic branching points (**Figure S2a”,a’’’**), thus suggesting a reduced complexity of the dendritic tree upon *Nr2f1* deletion.

**Figure 2.**
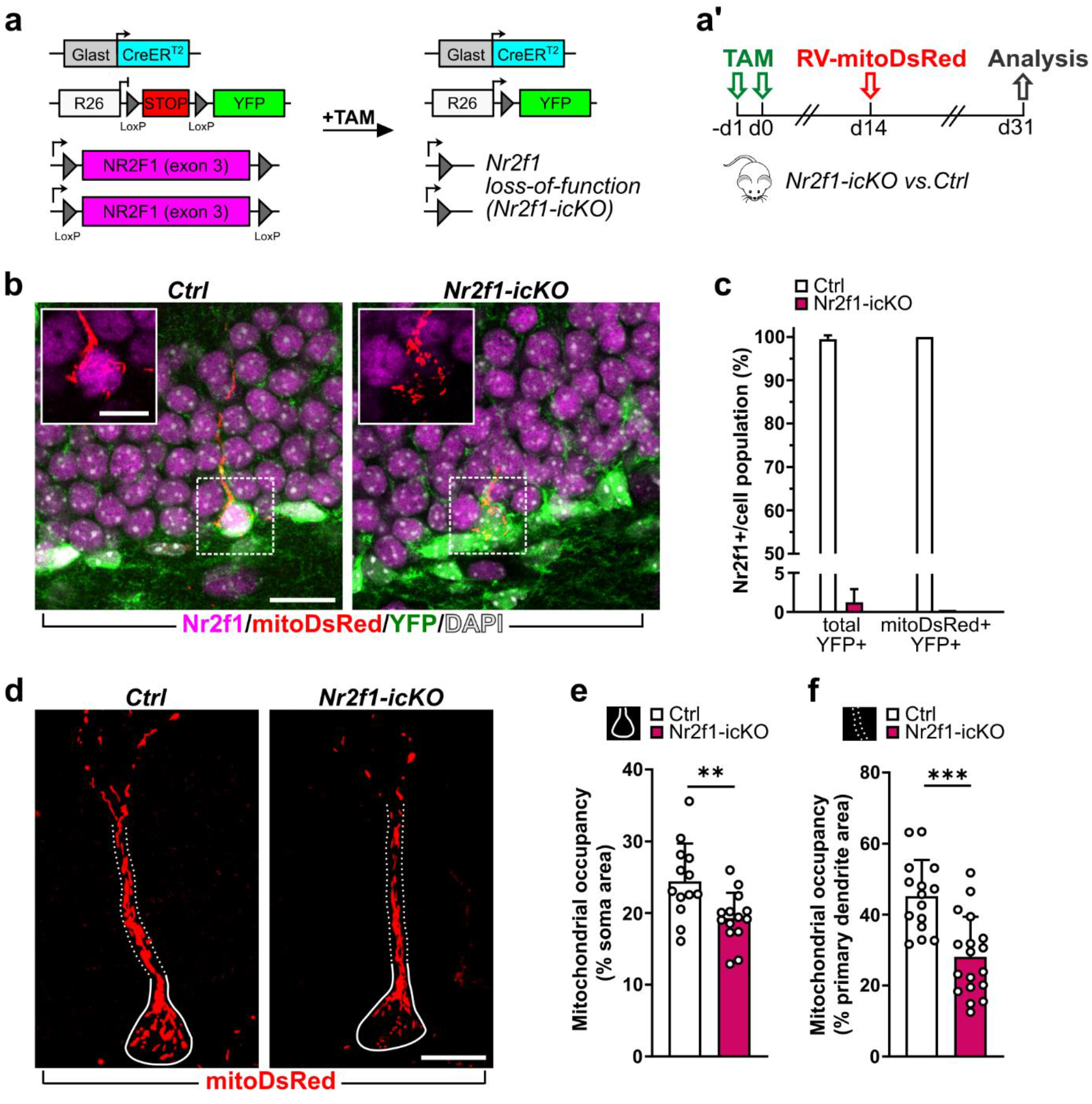
Nr2f1 LOF leads to decreased mitochondrial mass in the soma and proximal dendritic compartment of adult-born hippocampal neurons *in vivo*. **a.** Overview of the Cre-mediated gene rearrangements of Rosa26 (R26) and Nr2f1 loci in *Glast::CreERT2;R26-floxed STOP-Nr2f1fl/fl* mice. **a’.** Experimental design. **b.** Representative confocal images of adult-born neurons transduced with the RV-mitoDsRed (red) and immunostained for YFP (green) and Nr2f1 (magenta) in *Ctrl* and *Nr2f1-icKO* mice. Nuclei are counterstained with DAPI (white). **c.** Histogram reporting data validating Nr2f1 loss in the Glast-lineage. While in control DG, the vast majority of YFP+ and mitoDsRed+YFP+ cells showed nuclei that are positive for Nr2f1 (Nr2f1+), virtually all YFP+ and mitoDsRed+YFP+ cells in mutant DG are negative for Nr2f1. Nr2f1+ on total YFP+ cells: n=229/230 cells out of 3 *Ctrl* animals; N=5/340 cells out of 3 *Nr2f1-icKO* animals; Nr2f1+ on mitoDsRed+YFP+ cells: n=16/16 cells out of 3 *Ctrl* animals; n=0/20 cells out of 3 *Nr2f1-icKO* animals. **d.** Representative confocal images (max projection) showing the mitoDsRed+ mitochondria (red) within the soma area (solid lines) and primary dendrite (dotted lines) in control and *Nr2f1-icKO* mitoDsRed+DCX+YFP+ newborn neurons. **e,f.** Graph reporting the mitochondrial occupancy expressed as the percentage of the soma area (e) or of the primary dendrite area (f) covered by the mitoDsRed signal. Mann-Whitney test, p=0.0066 (e); p=0.0001 (f). For the analyses reported in e and f, n=13 (*Ctrl*) and 16 (*Nr2f1-icKO*) and n=14 (*Ctrl*) and 18 (*Nr2f1-icKO*) cells out of 3 animals per genotypes and at least 3 cells/animals were used. Data are shown as mean±SD. Each dot represents a cell. Scale bars: b, 20 μm; b (insets), 10 μm; d, 10 μm. *p<0.05, **p<0.01, ***p<0.001.

**Figure 3.**
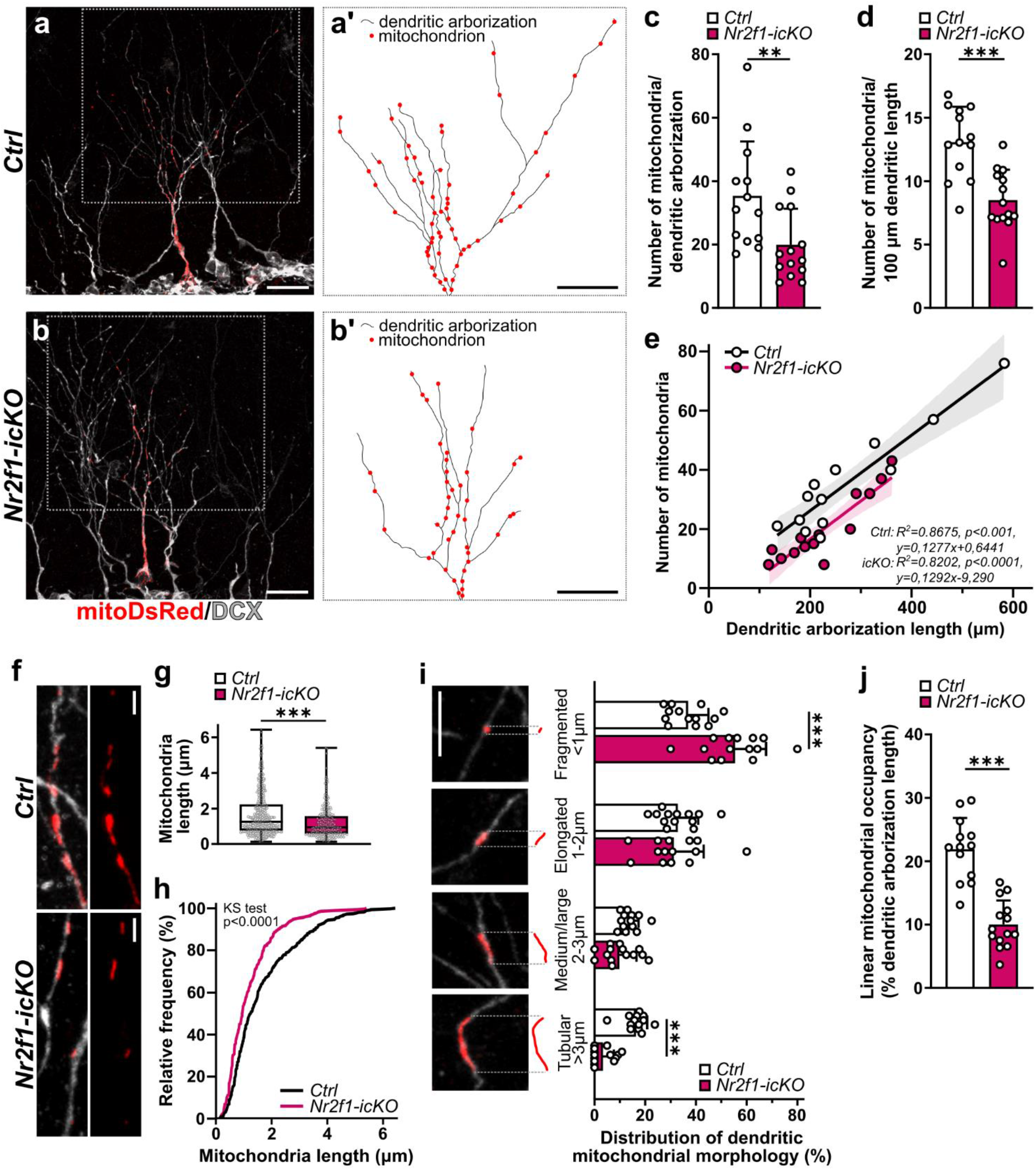
Alteration in dendritic mitochondrial number, architecture and mass in newborn neurons lacking Nr2f1. **a,b.** Representative confocal images of two recombined and mitoDsRed transduced (red) adult-born neurons immunolabelled with doublecortin (DCX, white) in the DG of both *Ctrl* (a) and *Nr2f1-icKO* (b). **a’,b’.** Representative drawing reporting the position of single mitochondrion (red dots) on the dendritic arborization of the two cells shown in a-b boxes. **c.** Histogram reporting the total number of mitochondria present in each dendritic arborization of the analyzed mitoDsRed+DCX+YFP+ cells. Mann-Whitney, p=0.0058. **d.** Histogram showing the linear density of dendritic mitochondria obtained by normalizing the total number of dendritic mitochondria to the total dendritic arborization length. Unpaired Student’s t test, p=0.0001. **e.** Graph showing the relationship between the dendritic arborization length (x axis) and the total number of dendritic mitochondria (y axis) per each mitoDsRed+DCX+YFP+ cell analyzed; linear regression with Pearson’s correlation was performed with R^2^, p-values and best fitted line equation displayed. Note the shift of the regression fitted line of *Nr2f1-icKO* cells without changes in its steepness compared to the *Ctrl* fitted line. **F.** Representative confocal images showing high magnification of the mitochondria in comparable level of dendrites in *Ctrl* and *Nr2f1-icKO* mitoDsRed+DCX+YFP+ adult-born neurons. **G.** Quantification of dendritic mitochondrial length in control and Nr2f1-depleted adult-born hippocampal neurons. Each data point represents an individual mitochondrial size; n=459 mitochondria out of 13 control cells, n=279 mitochondria out of 14 *Nr2f1-icKO* cells; Mann Whitney test, p<0.0001. **h.** Cumulative frequency distribution for the length of dendritic mitochondria in control vs. Nr2f1-ablated mitoDsRed+DCX+YFP+ adult-born neurons. Kolmogorov-Smirnov test, p<0.0001. **i.** Quantification of the distribution of dendritic mitochondrial morphologies into the four different subclasses based on their length (i.e., fragmented, <1 μm; elongated, from 1 to 2 μm; medium/large, from 2 to 3 μm; tubular, > 3 μm). Mann Whitney test: fragmented, p=0.001; elongated, p=0.6760; medium/large, p= 0.0917; tubular, p<0.0001. Each dot represents a cell. **j.** Graph reporting the mitochondrial occupancy expressed as the percentage of dendritic length covered by the mitoDsRed staining. Mann Whitney, p<0.0001. For all the analyses, n=13 (*Ctrl*) and n=14 (*Nr2f1-icKO*) cells out of 3 animals per genotype and at least 3 cells/animal were used for each analysis. Data are shown as: mean±SD (c,d,i,j); best regression fitted line in black, the band represent the 95% confidence interval and each dot represents a cell (e); box and whiskers plot with median (middle line), upper/lower quartiles, and error bars ranging from min to max values (g). Scale bars: a,b,a’,b’, 20 μm; f,i, 5 μm. *p<0.05, **p<0.01, ***p<0.001.

To initially assess the consequences of *Nr2f1* LOF on mitochondria in DG newborn neurons, we analyzed the mitochondrial occupancy expressed as the percentage of the area covered by the mitoDsRed+ signal over the area encompassing the perikaryon and the cone-shaped proximal portion of the primary dendrite (named “soma” hereafter) (**Figure 2d and Figure S2c**). MitoDsRed+ mitochondria appear densely packed and tightly organized into complex networks in the soma of control DCX+mitoDsRed+YFP+ newborn neurons (**Figure 2d,e and Figure S2b-b’’,c**). Interestingly, we observed a reduction by about 20% in the mitochondrial occupancy within the soma in Nr2f1-depleted neurons compared to control ones (**Figure 2d,e**). By exploiting the Mitochondrial Network Analysis (MiNA) toolset (Valente et al., 2017), we confirmed a decreased mitochondrial mass (i.e., mitochondrial footprint) and we further revealed a reduced complexity of the mitochondrial networks in Nr2f1-deficient newborn neurons (**Figure S2d**). In line, we found a reduction by 28% in the area covered by mitoDsRed+ mitochondria within the cylindrical portion of the primary dendrite of Nr2f1-deficient newborn neurons (**Figure 2d,f** and **Figure S2b’’’,e**).

We next evaluated the mitochondrial content and morphology in the distal dendrites of DCX+mitoDsRed+YFP+ newborn neurons, where high-resolution confocal microscopy allows the analysis of single mitoDsRed+ mitochondria (**Figure 3a-b’**) and thus enables quantification and in-depth morphometric assessment. We found a net reduction in the abundance of mitochondria in Nr2f1-depleted neurons not only when quantified as total number of mitochondria per dendritic arborization, but also when normalized to the total length of the dendritic arbor (**Figure 3c,d**). Even if the number of mitochondria linearly correlates to the dendritic arborization length both in *Ctrl* and in *Nr2f1-icKO* neurons (Spearman correlation: *Ctrl*, r=0.7868, p=0.0021; *Nr2f1-icKO*, r=0.8326, p<0.001), *Nr2f1-icKO* cells always show a lower number of mitochondria compared to control cells of akin dendritic length (**Figure 3e**), thus supporting that alteration in mitochondrial content is not secondary to morphological defects of the dendritic tree. On the other hand, previous studies indicate that mitochondria play a key role in the architecture of the dendritic arbor (Fang et al., 2016; Kimura and Murakami, 2014; Steib et al., 2014), thus altered dendritic mitochondria in newborn neurons lacking Nr2f1 might influence dendritic length and branching. Furthermore, we observed a reduction in the maximum length of single dendritic mitochondria in *Nr2f1-icKO* neurons compared to control ones (**Figure 3f-h**). By classifying mitochondria into four different categories based on their length [adapted from (Khacho et al., 2016)], we observed an increase in fragmented mitochondria concomitantly with a reduction in tubular ones in the dendrites of *Nr2f1-icKO* neurons (**Figure 3i**). In line, estimation of the linear mitochondrial occupancy (i.e., the percentage of the dendritic length covered by the mitoDsRed+ signal) unveiled a drastic decrease (about 55%) in Nr2f1-depleted neurons compared to controls (**Figure 3j**).

Overall, our morphometric analyses on mitoDsRed+ mitochondria revealed that loss of Nr2f1 function in adult-born hippocampal neurons induces a global impairment in the mitochondrial mass that appears progressively more dramatic from the proximal to the distal cell compartments (**Figure S2f**).

We finally exploited a complementary genetic strategy to increase *Nr2f1* expression (i.e. gain-of-function, GOF) in adult-born hippocampal neurons, by crossing the transgenic *CAGGS-lox-stop-lox-hCOUP-TFI* mouse line (Wu et al., 2010) to the *Glast::CreERT2* and *YFP* reporter line (herein *Nr2f1-O/E;* **Figure S3a**) (Bonzano et al., 2018). Analysis of the mitochondrial content on the dendritic arborization of DCX+mitoDsRed+YFP+ newborn neurons in *Nr2f1-O/E* mice revealed the occurrence of longer mitochondria without changes in their number, leading ultimately to an overall increase in the linear dendritic mitochondrial occupancy in newborn granule neurons (**Figure S3b-d**).

Data obtained by *Nr2f1* LOF and GOF, together with GO annotations disclosing a stronger enrichment for nuclear-encoded mitochondrial genes than that for other structural genes (e.g., cytoskeleton-related genes)-as Nr2f1 direct targets (**Figure 1**), strongly support a primary effect of Nr2f1 manipulation on mitochondria in adult-born DG neurons.

### The mitochondrial fusion factor Mfn2 is downregulated in mouse models of BBSOAS

Mitochondria undergo continuous morphological transition by dynamic processes thanks to the coordinated action of proteins promoting mitochondrial fission (e.g., Drp1, Fis1) and fusion (e.g., Mfn1/2, Opa1), as well as their transport (e.g., Miro, Milton) (Course and Wang, 2016; Giacomello et al., 2020; Misgeld and Schwarz, 2017). It is well known that altered expression and function of those factors leads to permanent changes in the overall shape of mitochondria (Chan, 2020) that could underlie the observed mitochondrial phenotype following *Nr2f1* LOF and GOF. We thus wondered whether altered Nr2f1 activity might impinge on mitochondrial dynamics. To this purpose, we exploited our ChIP-seq dataset for Nr2f1 (**Figure 1**) and unveiled that three of the key nuclear genes encoding proteins crucial for mitochondrial dynamics, namely dynamin-2 (Dnm2), mitochondrial dynamics protein of 51kDa (MiD5) and mitofusin-2 (Mfn2), are indeed bound by Nr2f1 on their promoters (**Figure 4a**). Based on the fragmented and elongated mitochondrial shape observed in *Nr2f1-icKO* and *Nr2f1-O/E* neurons respectively, we focused our attention on the fusion factor *Mfn2* as a putative target for Nr2f1 action. Notably, Nr2f1 binds to a promoter region closed to TSS of the *Mfn2* gene, which is associated to the H3K4me3 in the adult neural tissue (**Figure 4a**). We next crossed our Chip-seq dataset with previously published transcriptome data on the hippocampi of adult *Nr2f1-heterozygous* mice (*Nr2f1-het*) (Chen et al., 2020), a validated BBSOAS mouse model (Bertacchi et al., 2019a; Jurkute et al., 2021; Tocco et al., 2021). By Gene Set Enrichment Analysis (GSEA), we found that among the 1053 enriched genes approximately the 8% (82/1053) codes for mitochondrial proteins, including the *Mfn2* gene (**Figure 4b**). In addition, we did observe a reduction in Mfn2+ labelling (as the area occupied by the Mfn2+ signal and its intensity) in the DG of *Nr2f1-het* mice by *in situ* densitometric analysis (**Figure 4c,d,e**). To further confirm Mfn2 downregulation upon *Nr2f1* LOF, we quantified Mfn2 fluorescence labelling within the soma of triple-labeled Mfn2+DCX+YFP+ newborn neurons one months after TAM administration in *Nr2f1-icKO* mice (**Figure 4f,g**). In line with the observed reduction in mitochondrial occupancy in *Nr2f1-icKO* newborn neurons (**Figure 2e**), we found a 15% reduction in the percentage of area occupied by the Mfn2+ signal compared to controls (**Figure 4h,i**). Notably, analysis of the intensity of Mfn2 staining normalized to the area occupied by Mfn2+ signal (CTMF; about 40% decrease; **Figure 4j,k**), revealed a drastic drop, indicating that Nr2f1 deletion lowers Mfn2 levels in a cell-autonomous manner.

**Figure 4.**
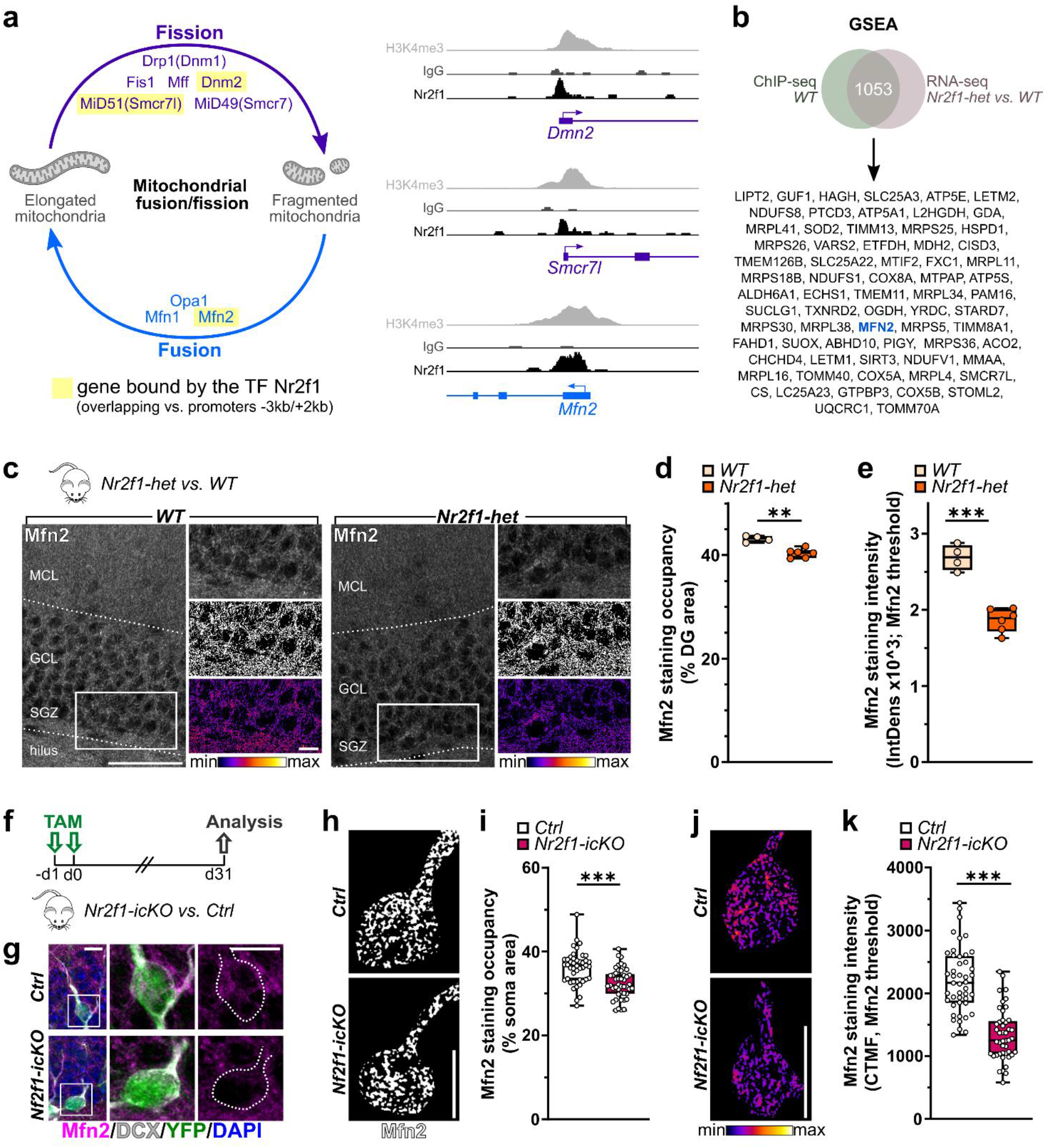
The fusion factor Mfn2 as a direct target for Nr2f1 transcriptional activity. **a.** Schematic illustration showing the main factors involved in the mitochondrial dynamics (fission in orange; fusion in light blue) in mammalian cells: among them, those underlined in yellow show specific Nr2f1 binding on their promoter regions as revealed by peak calling analysis on ChIP-seq data. Sequence tag accumulation of Nr2f1 ChIP-seq of around the TSS regions of Dmn2, Smcr7l and Mfn2 are shown. **b.** Gene Set Enrichment Analysis (GSEA) revealed Mfn2 among the genes encoding mitochondrial proteins bound by Nr2f1 and deregulated in constitutive heterozygous (*Nr2f1-het*) hippocampi. **c.** Representative confocal images showing Mfn2 staining within the DG of adult wildtype (*WT*) and *Nr2f1-het* mice. Insets show higher magnifications, the binarized images used to calculate the area occupied by Mfn2 staining and heat-map images of Mfn2+ signal in WT and Nr2f1-het DG. **d,e.** Graphs reporting the area occupied by Mfn2+ signal (d) and the intensity of Mfn2 protein staining (e) in *WT* and *Nr2f1-het* DG. Mann Whitney test: p=0.0095 (d) and unpaired Student’s t-test p<0.0001 (e). N=4 *WT* and 6 *Nr2f1-het* mice. **f.** Experimental design. **g.** Representative confocal images of the DG immunostained for YFP (green), Mfn2 (magenta), DCX (white) and counterstained with DAPI (blue) in *Ctrl* (top) and *Nr2f1-icKO* (bottom) mice one months after tamoxifen administration. **h.** Representative binarized images showing the area occupied by Mfn2 staining within the soma area in one control (top) and one Nr2f1-depleted (bottom) recombined newborn neurons (i.e. DCX+YFP+). **i.** Graph reporting the mitochondrial Mfn2 occupancy expressed as the percentage of the soma area covered by the Mfn2+ signal; Mann-Whitney test, p <0.0001; n=45 cells out of 3 Ctrl mice and n=45 cells out of 3 *Nr2f1-icKO* mice. **j.** Representative heat-map images of Mfn2+ signal in control (top) and Nr2f1-deficient (bottom) recombined newborn neurons. **k.** Graph reporting the mitochondrial Mfn2 staining intensity normalized on Mfn2 area residing in the cell compartment comprising the soma area (expressed as CTMF, cell threshold mitochondrial fluorescence); Unpaired t-test, p<0.0001; n=45 cells out of 3 Ctrl mice and n=45cells out of 3 *Nr2f1-icKO* mice. Data are shown as box and whiskers plot with median (middle line), upper and lower quartile, and the error bars ranging from min to max values. Each dot represents an animal (d,e) or a cell (i,k). c, 50 μm (low magnification) and 10 μm (higher magnification). Scale bars: g,h,j, 10 μm; ***p<0.0001.

It is worth noting that the morphological changes we did observe in adult-born neurons lacking Nr2f1, including mitochondrial fragmentation and decreased mitochondrial mass, as well as a progressively stronger phenotype proceeding from proximal to distal cell compartments (**Figure S2f**), recapitulate the phenotype of Mfn2 downregulation described in differentiated excitatory neurons *in vitro* (Fang et al., 2016). Overall, it is thus highly conceivable that Nr2f1 might act as upstream regulator of Mfn2 expression controlling mitochondrial dynamics in adult-born DG neurons.

## Conclusion

Mitochondria, which exert essential roles in both bioenergetic and non-energetic biological processes, are increasingly recognized as major players in neuronal development and function, including neural stem cell activation, dendritogenesis, synaptic transmission and plasticity (Faits et al., 2016; Khacho et al., 2019, 2016; Kimura and Murakami, 2014; Rangaraju et al., 2019b, 2019a; Steib et al., 2014). Accordingly, mitochondrial dysfunction dramatically contributes to the pathogenesis of various neurodegenerative and neurodevelopmental disorders (Jurcau, 2021; Monzio Compagnoni et al., 2020; Rojas-Charry et al., 2021). Here, by exploiting genetic mouse models we clearly demonstrated the occurrence of a mitochondrial phenotype in neurons lacking *Nr2f1* function, including altered mitochondrial mass and shape. As the homology between human and mouse Nr2f1 is very high (especially in the DNA binding domain with a 100% of amino acid sequence homology) (Alfano and Studer, 2013; Qiu et al., 1995), their functions and targets are likely to be conserved in both species, strongly suggesting the implication of mitochondrial dysfunction in BBSOAS neuropathology. Accordingly, alterations in mitochondrial energy supply due to a defective function of the electron transport chain (ETC), have been reported in muscle biopsies from two BBSOAS patients (Hobbs et al., 2020; Martín-Hernández et al., 2018). Furthermore, in light of our findings, the mitochondrial abnormalities recently described in the optic nerve and retina of *Nr2f1* constitutive knockout/het mice (Bertacchi et al., 2019a) might be a direct consequence of Nr2f1 mutations and contribute to the optic atrophy in Nr2f1 mouse models.

Altogether, our data point to mitochondrial dysfunction in neural tissue as a key pathological mechanism in BBSOAS, suggesting that the current estimation of mitochondrial involvement in BBSOAS patients might be underestimated. Beside this rare disease, altered Nr2f1 expression has been reported in neural tissues and cells of Down syndrome (Bhattacharyya et al., 2009; Halevy et al., 2016) as well as neurodegenerative disorders, including mouse models of Alzheimer’s and Parkinson’s diseases (Walter et al., 2021; Zheng et al., 2020), thus our findings open new perspectives for the future investigation on the implication of Nr2f1 in the mitochondrial dysfunction associated with their pathogenesis and progression.

## Material and Methods

### Animals and treatments

*In vivo* experiments were performed on two to three-month-old C57BL/6J mice (Charles Rivers) of both genders. Triple transgenic mice (C57BL/6J background) were also used to manipulate Nr2f1 expression through tamoxifen administration. *Glast::CreERT2^+/wt^*;*R26-loxP-stop-loxP-YFP^yfp+/yfp+^*;*Nr2f1^fl/fl^* (*Nr2f1-icKO*) were used for *in vivo* loss-of-function experiments and *Glast::CreERT2^+/wt^*;*R26-loxP-stop-loxP-YFP^yfp+/yfp+^*;*CAG-S-loxP-stop-loxP-hCOUP-TFI^+/wt^* (*Nr2f1-O/E*) were used for *in vivo* Nr2f1 overexpression [see (Bonzano et al., 2018)]. *Glast::CreERT2^+/wt^*;*R26-loxP-stop-loxP-YFP^yfp+/yfp+^* were used as control mice (*Ctrl*). For activation of the *CreERT2*-recombinase in the Glast+ lineage, animals were administered with tamoxifen (TAM, SIGMA, T-5648) at the dose of 2,5 mg/mouse/day dissolved into corn-oil (SIGMA) by means of intraperitoneal injections (i.p.) for 2 consecutive days at the age of two-months (Bonzano et al., 2018). Adult (four-month-old) constitutive *Nr2f1* heterozygous mice (i.e. *Nrf1^wt/null^*, named *Nr2f1-het*) were generated and genotyped as previously described (Jurkute et al., 2021). Littermates of *Nr2f1-het* mice with wildtype Nr2f1 alleles were used as controls (i.e., *Nr2f1^wt/wt^*. WT). Both *Nr2f1-het* and WT littermates were bred in a 129S2/SvPas background. Mice were housed under standard laboratory conditions (n=2-4 mice/standard cage) with basic recommended environmental enrichment (paper tubes and litter, igloo) under a 12 h light/dark cycle with access to food and water *ad libitum*. All procedures were conducted in accordance with the Guide for the Care and Use of Laboratory Animals of the European Community Council Directives (2010/63/EU and 86/609/EEC) and approved by local bioethics committees, the Italian Ministry of Health (Authorization number 864/2018-PR, protocol E6B5E.3) and the French Ministry of Education, Research and Innovation CIEPAL NCE/2019–548 (Nice) under authorization #15 349 and #15 350.

### Retroviral production

Replication-deficient recombinant murine Moloney leukemia (MML) retroviruses specifically transduce proliferating cells and allow the dating of their birth, as well as label precursor cells and their progeny in the adult hippocampal neurogenic lineage (Zhao et al., 2006). CAG-IRES-mitochondrial Discosoma red (mitoDsRed) were generated from the pCAG IRES–GFP vector (Jagasia et al., 2009) by replacing the GFP coding sequence with cDNA for mitochondrially targeted DsRed. Retroviruses were generated as described previously(Steib et al., 2014; Zhao et al., 2006). Virus-containing supernatant was harvested four times every 48 h after transfection and concentrated by two rounds of ultracentrifugation. Viral titers were determined by transduction of HEK293T cells for 72 h with limiting dilution of MMLV suspension and counting of reporter expressing cell spots under a fluorescent microscope (Leica Microsystems). Titers applied were ~5 × 10^8^ colony-forming units (CFU)/ml (Zhao et al., 2006).

### Surgical procedure for retroviral injection

Two weeks after TAM administration, adult mice were anesthetized in a constant flow of Isofluorane (3%) in oxygen, positioned in a stereotaxic apparatus (Stoelting) and injected with a pneumatic pressure injection apparatus (Picospritzer II, General Valve Corporation). The skull was exposed by an incision in the scalp and a small hole (about 1 mm) drilled through the skull. 0.8 μl of retrovirus-mitoDsRed was injected in the DG using a sharpened glass capillary at the following stereotaxic coordinates: −2 mm (antero-posterior), 1.5 mm (lateral) to Bregma and −2.0 mm below the surface of the skull. Mice (n=3/4 genotype/experiments) were analyzed 17 days after retroviral injection.

### Tissue preparation and immunofluorescence staining

For immunostaining, adult mice were deeply anesthetized with an intraperitoneal injection of a mixture of tiletamine and zolazepam (40-80 mg/kg; Zoletil; Virbac) and perfused transcardially with ice-cold 0.9% saline solution followed by ice-cold 4% paraformaldehyde (PFA) in 0.1 M phosphate buffer (PB), pH 7.4. Brains were removed from the skull, post-fixed for 4 hours in the same PFA solution, cryoprotected in a 30% sucrose solution (in 0.1 M PB, pH 7.4), embedded with cryo-embedding matrix (OCT; Killik, BioOptica), frozen at −80°C and finally sectioned using cryostat (Leica Microsystems). Free-floating coronal serial sections (50μm-thick for the hemisphere ipsilateral to the viral injection; 40μm-thick for the contralateral hemisphere) were collected in series on multi-well dishes (6 to 8 wells per animal). Sections were stored at −20°C in antifreeze solution (30% ethylene glycol, 30% glycerol, 10% PB: 189 mM NaH_2_PO_4_, 192.5 mM NaOH; pH 7.4) until use. Immunofluorescence reactions were performed on free-floating coronal serial sections as detailed below: sections were incubated in blocking solution (0.01M PBS - pH 7.4, 1 or 2% Triton X-100 for 40 and 50 μm-thick slices respectively, and 10% normal serum of the same species of the secondary antibody, i.e., normal donkey serum – NDS) for two hours at room temperature. Afterwards, slices were incubated for 48 hours at 4°C with primary antibodies (see Table 1) diluted in 0.01M PBS (pH 7.4), 1% to 2% Triton X-100 (for 40 and 50 μm-thick slices respectively), and 1% NDS. For immunostainings against Mfn2, sections were subjected to antigen retrieval. Slices were incubated in 10mM Tris, 1mM EDTA, 0.05% Tween 20 for 2 min at 99°C and washed three times with MilliQ water followed by three washing step with 0.01M PBS (pH 7.4) prior incubation with primary antibody. Sections were washed in PBS and incubated for 2 hours or overnight (for 40 and 50 μm-thick slices respectively) at 4°C with secondary antibodies (see Table 2) in 0.01 M PBS (pH 7.4), 0.2% Triton X-100, and NDS (1%). Sections were washed in 0.01M PBS (pH 7.4) and incubated for 20 minutes at room temperature with 4,6-diamidino-2-phenylindole (DAPI, 1 μg/ml) to label nuclei. Sections were washed in 0.01M PBS (pH 7.4) then mounted on gelatine coated slides, air dried, and coverslipped with antifade mounting medium Mowiol (4-88 reagent, Calbiochem 475904).

**Table 1.** List of the primary antibodies.

| Antigen name | Host | Dilution | Source | Catalogue number |
| --- | --- | --- | --- | --- |
| **Primary antibodies** |  |  |  |  |
| COUP-TFI/Nr2f1 | Rabbit | 1:1000 | Abcam | ab181137 |
| COUP-TFI/Nr2f1 | Rabbit | 1:1000 | Studer's lab | - |
| DCX | Goat | 1:1500 | Santa Cruz Biotechnology | Sc-8066 |
| GFAP | Goat | 1:2000 | Abcam | ab53554 |
| GFP | Chicken | 1:1000 | AvesLab | GFP-1020 |
| Mfn2 | Rabbit | 1:500 | Immunological Science | MAB-94608 |
| NeuN | Mouse | 1:1000 | Chemicon | MAB377 |

**Table 2.**
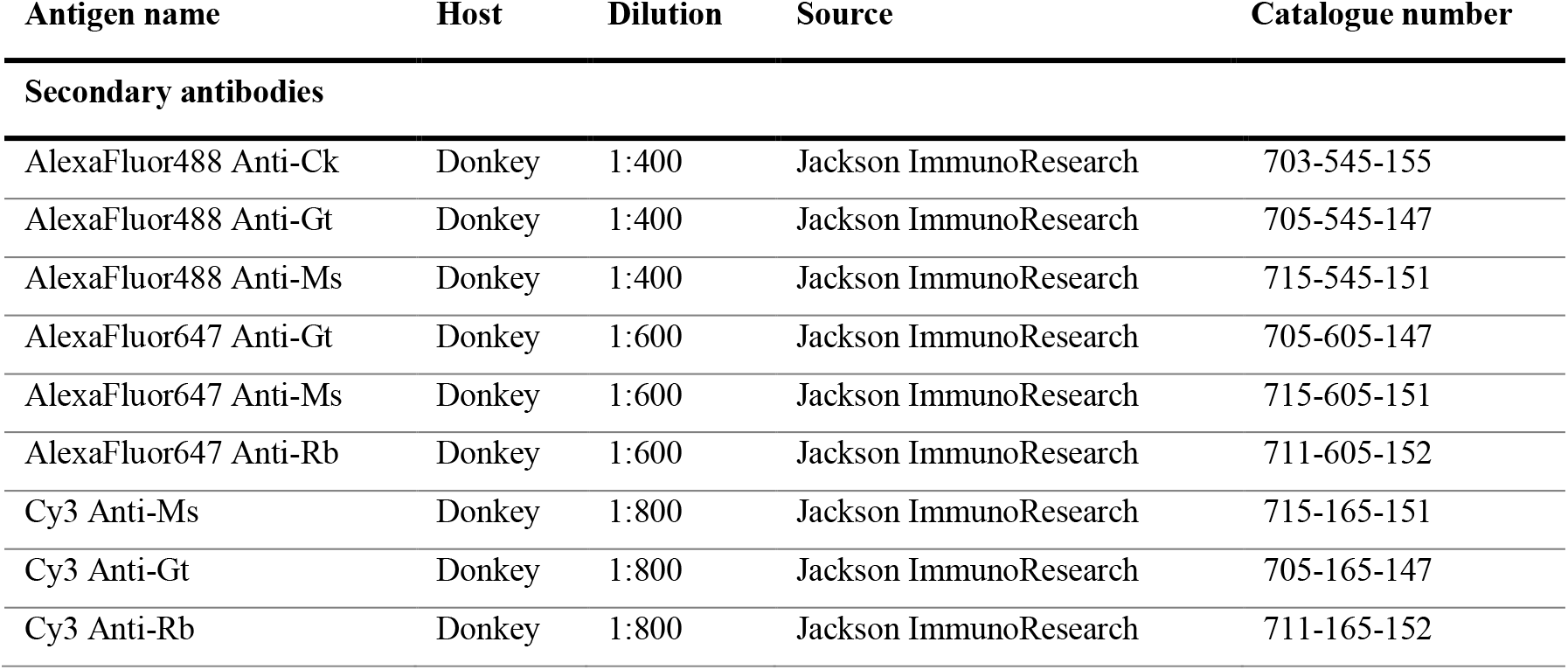
List of the secondary antibodies.

| Antigen name | Host | Dilution | Source | Catalogue number |
| --- | --- | --- | --- | --- |
| **Secondary antibodies** |  |  |  |  |
| AlexaFluor488 Anti-Ck | Donkey | 1:400 | Jackson ImmunoResearch | 703-545-155 |
| AlexaFluor488 Anti-Gt | Donkey | 1:400 | Jackson ImmunoResearch | 705-545-147 |
| AlexaFluor488 Anti-Ms | Donkey | 1:400 | Jackson ImmunoResearch | 715-545-151 |
| AlexaFluor647 Anti-Gt | Donkey | 1:600 | Jackson ImmunoResearch | 705-605-147 |
| AlexaFluor647 Anti-Ms | Donkey | 1:600 | Jackson ImmunoResearch | 715-605-151 |
| AlexaFluor647 Anti-Rb | Donkey | 1:600 | Jackson ImmunoResearch | 711-605-152 |
| Cy3 Anti-Ms | Donkey | 1:800 | Jackson ImmunoResearch | 715-165-151 |
| Cy3 Anti-Gt | Donkey | 1:800 | Jackson ImmunoResearch | 705-165-147 |
| Cy3 Anti-Rb | Donkey | 1:800 | Jackson ImmunoResearch | 711-165-152 |

### Microscope acquisition and quantifications

Images of multiple immunofluorescence on tissue sections were acquired with a TCS SP5 confocal microscope (Leica Microsystems) or a Nikon microscope coupled with a computer-assisted image analysis system (Neurolucida software, MicroBrightField). Confocal z-stacked images used for morphometric analyses on both cellular and mitochondrial architecture were captured through the thickness of the slice (50 μm) at 0.5-μm optical steps with an objective 63X/1.4 N.A. oil immersion lens with an additional zoom (2x) and a resolution of 8-bit, 1024/1024 pixels and 50Hz scan-speed (1 voxel=133.6*133.6*395.2 nm, xyz). mitoDsRed+YFP+ newborn granule neurons residing in the upper and lower blades of the DG were used for analyses only when satisfying all the following criteria: (i) to be positive for doublecortin (i.e. triple DCX+mitoDsRed+YFP+ newborn cells); (ii) to bear an apical dendrite that arborize into the molecular cell layer (MCL); (iii) to show a high and homogeneous mitoDsRed signal; and (iv) to include the whole dendritic morphology within the 50-μm section thickness. Morphometric analyses of the selected DCX+mitoDsRed+YFP+ newborn neurons was performed by using the Simple Neurite Tracer (SNT of the Neuroanatomy toolkit) plug-in on Fiji (https://fiji.sc/). Perikarya size was assessed by manually drawing the largest cross-sectional area of each cell soma based on the DCX+YFP+ double staining using the polygon selection tab in Fiji. Manual counting of dendritic mitochondria number and evaluation of their length was carried out on 2D images obtained by the Max intensity projection tool applied on the optical slices that include the whole dendritic arborization. The number of dendritic mitochondria was manually analyzed by the Cell Counter and channel tool plug-ins; measurement of the dendritic mitochondrial length was carried out by drawing the length (parallel to the dendrite) of each mitochondrion using the segmented line tool on Fiji.

Analyses of the mitochondrial mass in the soma and the primary dendrite compartments were carried out as described below. First, for each cell compartments a maximum z-projection of all the optical slices including the region of interest (ROI) was applied and the ROI based on the cytoplasmic YFP+ signal on Fiji, drawn and measured as area in μm^2^. In particular, the soma area comprises the perikaryon area *plus* the cone-shaped portion of the primary dendrite directly stemming from the soma, whereas the area of the primary dendrite includes the cylindric part of primary apical dendrite to its first ramification (see **Figure S2b**). Afterwards the channel containing the mitochondrial signal (i.e., mitoDsRed) was binarized after thresholding and the resulting binary image was used to calculate the area covered by mitochondrial signal and expressed as a percentage on the ROI area. In addition, the MiNA toolkit developed by the StuartLab (Valente et al., 2017) was used for the semi-automated analyses of the mitochondrial network features in the soma compartment.

To quantify Mfn2 immunostaining within the soma of DCX+YFP+ neurons, confocal image z-stacks were captured throughout the thickness of the cell bodies of DCX+YFP+ neurons. Acquisitions were done with 0.5-μm optical step size using an objective 63X/1.4 N.A. oil immersion lens and by adding an additional zoom (3x) with a resolution of 8-bit, 1024/1024 pixels and 50Hz scan-speed (1 voxel=120.3*120.3*395.2 xyz). Importantly, all the parameters (i.e., laser power, gain and offset) were kept the same among acquisitions. We developed a custom-made workflow allowing an unbiased and semi-automated quantitative analysis of Mfn2 staining coverage and intensity on triple labeled Mfn2+DCX+YFP+ neurons on Fiji. First, a max z-projection of the optical slices containing the whole soma area was applied and the soma ROI was manually drawn based on the cytoplasmic YFP+ signal. The obtained ROI was applied to the channel including the naïve Mfn2 signal to calculate the Mfn2 staining intensity within the soma shaft compartment and expressed as the Corrected Total Cell Fluorescence (CTCF) calculated by the formula CTCF = Integrated Density – (ROI X Mean fluorescence of background readings). The background readings were calculated by averaging three mean fluorescence obtained by three small ROI (5×5μm) without Mfn2 staining (i.e., blood vessel lumen or cell nuclei close to the cell of interest). To evaluate the fractional area covered by Mfn2 staining in the soma area, we first applied a “pre-processing” step on the acquired channel including the Mfn2 signal as follow: (i) unsharp mask (radius=2, mask=0.4); (ii) substract background (rolling=50); (iii) enhance local contrast (i.e., CLAHE; blocksize=9, histogram=256, maximum=4, no mask; slow); (iv) median filtering (radius=1). Afterwards, the channel containing the Mfn2 staining was given a threshold (by Moments algorithm) and the binary image was cleaned outside the ROI to get the area occupied by Mfn2 staining (i.e., Mfn2 ROI) within the selected cell. Finally, we calculated the area fraction covered by Mfn2 signal on the soma area (i.e., Mfn2 staining occupancy) and expressed it as a percentage. The Mfn2 ROI was also applied to the naive channel containing the Mfn2 signal to calculate the Mfn2 staining intensity both as Integrated Density (IntDens) and as corrected total mitochondrial fluorescence (CTMF) wherein the ROI was the Mfn2 signal or calculated as the CTCF wherein the ROI was the soma area. To calculate the Mfn2 staining coverage on the DG of adult constitutive *Nr2f1-het* mice, we adopted the same strategy descripted above but on the complete image. For all the analyses the number of cells, animals, and mitochondria are described in the figure legends.

### Chromatin immunoprecipitation followed by sequencing (ChIP-seq)

Adult neocortical tissue was acutely isolated from two-month-old C57BL/6J mice (n=3 males). Animals were deeply anesthetized with an intraperitoneal injection of a mixture of tiletamine and zolazepam (40 mg/kg; Zoletil; Virbac) and xylazine (5 mg/kg; Rompun; Bayer) and the brain removed after rapid decapitation. Neocortices from both hemispheres were microdissected and pooled together, frozen and stored at −80°C until further analysis. Around 10 mg of neural tissue were chopped with a scalpel, harvested in 5 ml of 0.01 M PBS (pH 7.4) and crosslinked by adding formaldehyde to a final concentration of 1.5%, incubated for 30 min at RT on a rotator, and quenched with 0.125 M glycine for 5 min at RT. After crosslinking, the tissue was washed twice in cold 0.01 M PBS (pH 7.4) and centrifuged at 1.000×g for 5 min at 4°C. Tissue pellet was resuspended in 0.25ml of sodium dodecyl sulfate (SDS) lysis buffer (50 mM Tris pH 8.0, 1% SDS, 10 mM EDTA, anti-proteases) and incubated on a rotator for 30 min at 4°C. Afterwards, sample was sonicated for 18 cycles on high power setting (30s ON, 30s OFF) using the Bioruptor Next Gen (Diagenode) and then centrifuged at 20.000xg for 10 min at 4°C.

The isolated chromatin was diluted 10-fold with ChIP dilution buffer (16.7 mM Tris-HCl pH 8.0, 0.01% SDS, 1.1 % Triton X-100, 1.2 mM EDTA, 167 mM NaCl) (1/10 was kept as input) and incubated with 2 μg of rabbit anti-COUP-TFI/Nr2f1 antibody from Studer’s lab (Tripodi et al., 2004) overnight at 4°C on a rotator. Protein G-conjugated magnetic beads (Dynal, Thermo Fisher Scientific) were saturated with PBS/1% BSA overnight at 4°C. Next day, samples were incubated with saturated beads for two hours at 4°C on a rotator, and subsequently washed with 1 ml of cold Low salt buffer (20 mM Tris-HCl pH 8.0, 0.1 % SDS, 1% Triton X-100, 2 mM EDTA, 150 mM NaCl), 1 ml of cold High salt buffer (20 mM Tris-HCl pH 8.0, 0.1 % SDS, 1% Triton X-100, 2 mM EDTA, 500 mM NaCl), 1 ml of cold LiCl buffer (10 mM Tris-HCl pH 8.0, 1% DOC, 250 mM LiCl, 1 mM EDTA, 1% NP-40), and twice with 1 ml of cold TE buffer (10 mM Tris-HCl pH 8.0, 1 mM EDTA). The immunoprecipitated chromatin was eluted with 200 μl of Elution buffer (10 mM Tris-HCl pH 8.0, 1 mM EDTA, 1% SDS, 150 mM NaCl, 5mM DTT) for 30 min at RT on a rotator, and decrosslinked at 65°C overnight. The decrosslinked DNA was purified using QIAquick PCR Purification Kit (Qiagen) according to the manufacture’s instruction.

For genome-wide analysis of binding, sequencing libraries were constructed using the NEBNext ChIP-seq Library Prep Reagent Set for Illumina and a NextSeq 500 Illumina sequencer (New England BioLabs Inc.).

### Bioinformatic analyses

The reads from sequencing were mapped to the mouse genome (mm9 assembly) using Bowtie version 0.12.7, reporting only unique hits with up to two mismatches. The redundant reads were collapsed, and peak calling was performed using MACS version 1.4.1 (Zhang et al., 2008) with normalization for IgG ChIP at a fixed p-value=1E-8. Homer version 4.11 was used to perform de-novo motif discovery and peak annotation. A ChIP-seq peak was considered associated with a gene if it showed an overlap of −3kb/+2kb region around a transcription start site (TSS) of the gene. Gene Ontology (GO) enrichment analysis was performed using PANTHER Overrepresentation Test version 14 (Mi et al., 2019).

We performed a pre-rank Gene Set Enrichment Analysis (GSEA) using as “rank value” the DeSeq2 statistic of genes expression in the hippocampi of adult *Nr2f1-het vs*. WT mice obtained by bulk RNA-seq by Chen and colleagues (Chen et al., 2020), and as “pathway” the genes bound on the promoter by Nr2f1.

### Statistics

Data are derived from at least three different animals/group. A two-tailed unpaired t-test was performed to compare the differences between two groups, moreover, an F-test of equality of variances was conducted to compare variances, and Welch’s correction was applied in case of unequal variance distribution. Data distribution was tested to be normal with either the Shapiro-Wilk test (alpha=0.05; if 3<n<50) or D’Agostino-Pearson (if n>50). When comparing two populations of data, two-tailed unpaired Student’s t test was used to calculate statistical significance in cases of Gaussian distribution; otherwise, the non-parametric Mann-Whitney test was used. For Sholl analysis, two-way repeated measures ANOVA with Sidak’s post-hoc test was used. The confidence interval was expressed with 95% confidence. For the analysis of normalized data (i.e., fold change) a Kruskal-Wallis test followed by Dunn’s post-hoc test was used. Kolmogorov-Smirnov test was carried out to compare cumulative distributions. The statistical significance was defined as follows: *p<0.05, **p<0.01, and ***p<0.001. All statistical analyses were performed using Graphpad Prism 8 software (version 8.0.2; GraphPad Software, San Diego California USA, www.graphpad.com). Fisher’s exact test, with the Benjamini–Hochberg False Discovery Rate (FDR) correction for multiple testing, has been added as the default algorithm for the over-representation test for GO analyses (Mi et al., 2019).

## Acknowledgements

This work received the financial support of the Jerome Lejeune Foundation (Cycle 2020a – Project #1977) and was supported by the ex60% (University of Turin) to S. De Marchis and the Foundation Cassa Risparmio Torino (CRT – 2019/2254) to S. Bovetti. Postdoctoral fellowships to S. Bonzano were funded by: Compagnia San Paolo (Turin) “Bando per l’internazionalizzazione della ricerca”; Fondazione Umberto Veronesi (Milan); the Accademia Nazionale dei Lincei “Centro Linceo Interdisciplinare Beniamino Segre” (Rome).

## Author Information

The authors declare no competing financial interests.

## Supplemental Figures with Legends: S1, S2, S3

**Figure S1 - Related to Figure 1. a,b.**
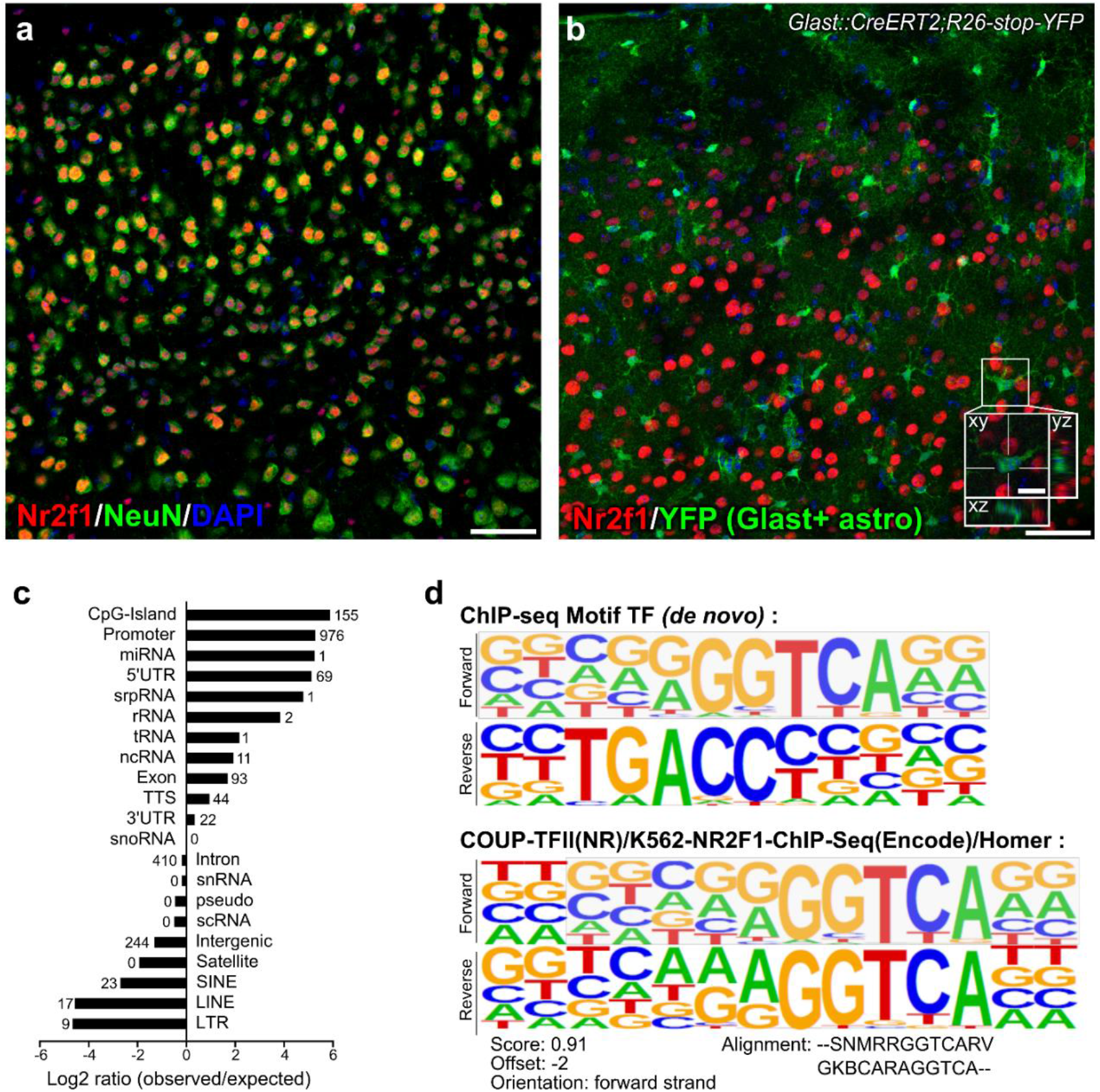
Representative confocal images showing Nr2f1 immunofluorescence in the neuronal population within the adult mouse neocortex., The vast majority of Nr2f1+ cells belongs to the neuronal population, as revealed by the high colocalization of Nr2f1 immunofluorescence with the neuronal marker NeuN (a) and no co-expression within the recombined astrocytic population labelled by YFP in *Glast::CreERT2;Rosa26-lox-stop-lox-YFP* mice following a chase of two weeks after tamoxifen administration (b). **c.** Graph reporting the logarithmic ratio between the observed genomic Nr2f1-bound peaks versus the expected ones. Numbers in each line represent the quantity of annotated peaks/annotation category. **d.** Raw data showing the *de novo* motif revealed by ChIP-seq for Nr2f1 in the adult brain and one of the best matches with known ChIP-seq database. Scale bars: a,b, 50 μm (low magnification); b, 10 μm (higher magnification with resliced).

**Figure S2 - Related to Figure 2 and 3.**
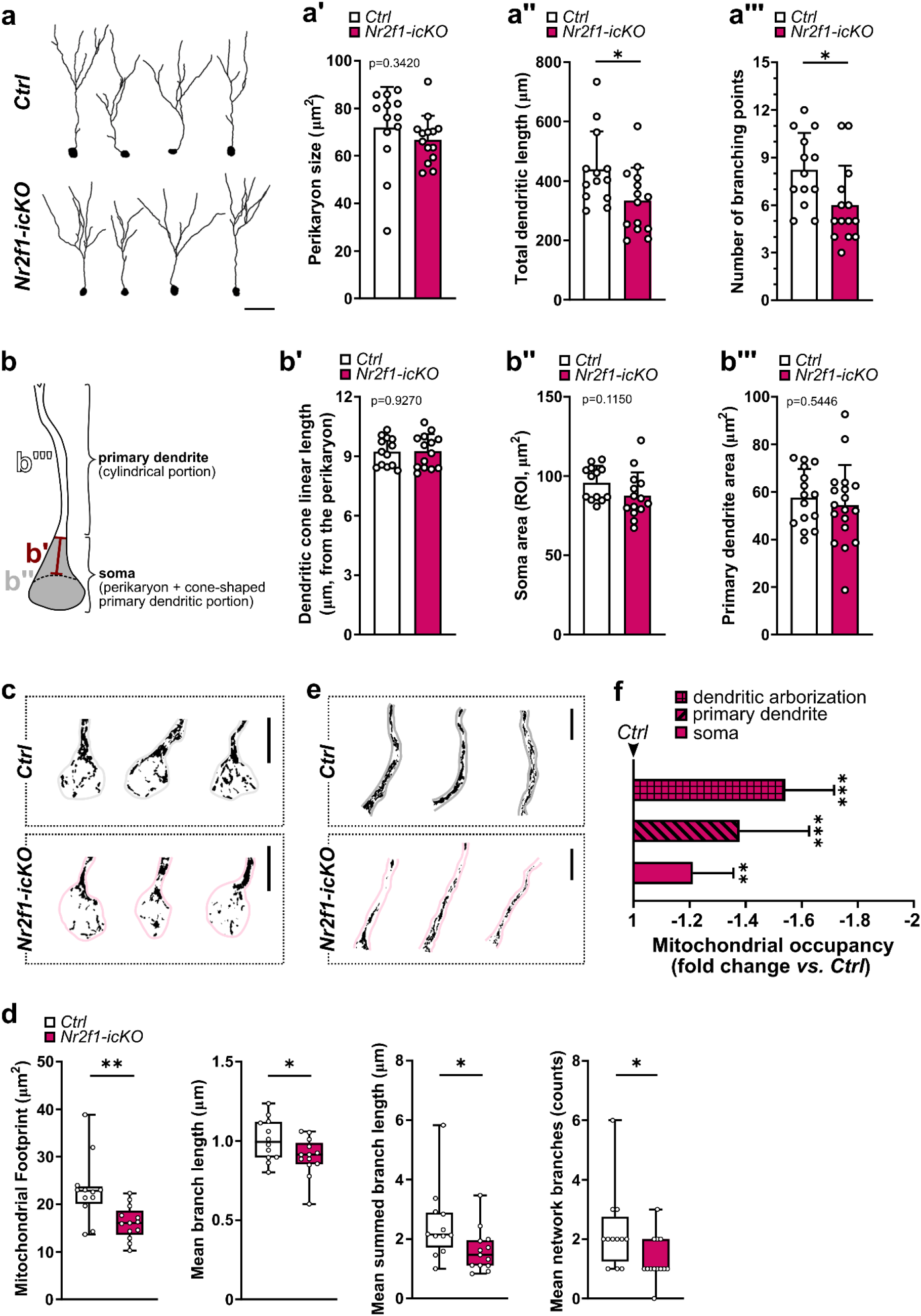
**a.** Representative reconstruction of triple labelled mitoDsRed+DCX+YFP+ DG newborn neurons at 17dpi and used for the analyses. **a’.** Histogram reporting no difference in the perikarya of control vs. Nr2f1-deficient mitoDsRed+DCX+YFP+ newborn neurons. Unpaired Student’s t-test. **a’’,a’’’.** Histograms showing the total dendritic length (a’’; unpaired Student’s t-test p=0.0296) and the number of dendritic branches (a’’’; Mann-Whitney t-test p=0.0153) in Nr2f1 deficient mitoDsRed+DCX+YFP+ newborn neurons and control neurons. For all the analyses in a’ -a’’’, n=13 and n=14 cells out of 3 animals per genotypes and at least 3 cells/animals were used for each analysis. **b.** Drawing showing the ROIs used for the quantifications of mitochondrial occupancy and quantified in b’, b’’ and b’’’. **b’-b’’’.** Quantification of the morphometric parameters used to calculate the mitochondrial occupancy. Student’s unpaired t test, exact p-values in the graphs. n=13 *Ctrl* and n=14 *Nr2f1-icKO* cells out of 3 animals per genotype and at least 4 cells/animal. Data are shown as mean±SD. **c.** Representative binarized images showing the mitoDsRed+ mitochondria (black) within the soma (c) in control and Nr2f1-deficient mitoDsRed+DCX+YFP+ newborn neurons. **d.** Graphs showing the results obtained by applying the MiNA toolkit on the mitoDsRed signal within the soma compartment of control and *Nr2f1-icKO* newborn mitoDsRed+DCX+YFP+ neurons. From the left to the right: unpaired Student’s t-test with Welch’s correction, p=0.0061; unpaired Student’s t-test, p=0.0465; Mann-Whitney test, p=0.0248; Mann-Whitney test, p=0.0341. Data are shown as box and whiskers plot with median (middle line), upper and lower quartile, and the error bars ranging from min to max values. A coherent reduction in the parameter of mitochondrial footprint (i.e. the total area consumed by mitochondrial signal after being separated from the background) with a coincident decrease in the length of branches (i.e. mean length of all the lines used to represent the mitochondrial structures) and in the summed branch length (i.e. the mean of the sum of the lengths of branches for each independent mitochondrial structure divided by the number of independent mitochondrial structures) as well as in the number of network branches (i.e. the mean number of attached lines used to represent each mitochondrial structure) was found in *Nr2f1-icKO* neurons compared to controls. **e.** Representative binarized images showing the mitoDsRed+ mitochondria (black) within the primary dendrite in control and Nr2f1 -deficient mitoDsRed+DCX+YFP+ newborn neurons. **f.** Histogram showing the reduction (expressed as fold change) in the mitochondrial mass within the different subcellular compartments in Nr2f1 -depleted mitoDdRed+DCX+YFP+ cells compared to the one measured in the compartments of the control cells. Statistics (asterisks) were obtained by Mann Whitney test of fold change values obtained from Nr2f1-icKO cells and normalized to control values (Nr2f1-icKO vs. Ctrl): soma, p=0.0047; primary dendrite, p=0.0002; dendritic arborization, p<0.0001. Each dot represents a cell. Scale bars: a, 30 μm; c,e, 10 μm. *p<0.05, **p<0.01, ***p<0.001.

**Figure S3.**
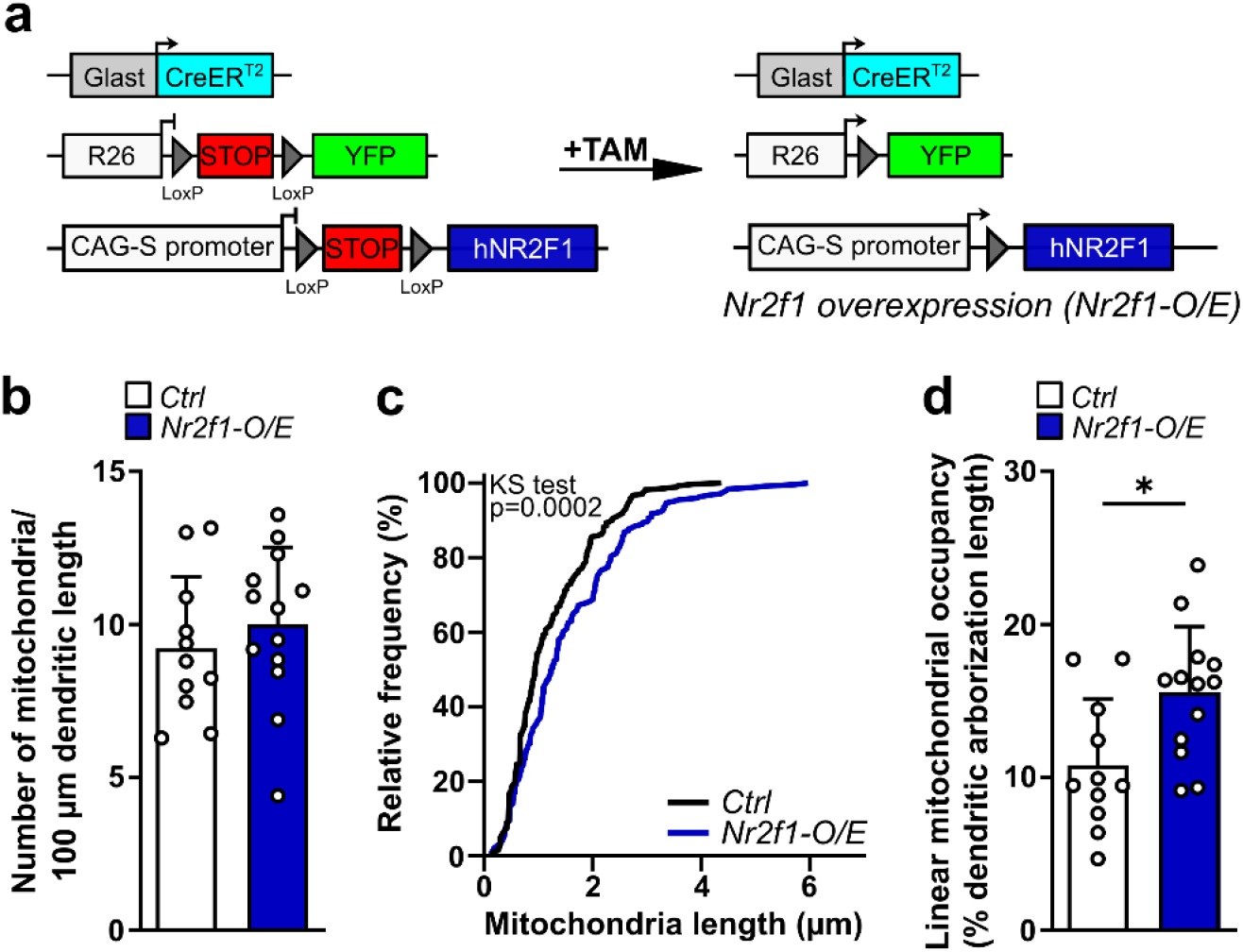
Nr2f1 overexpression leads to increased mitochondrial content in adult-born DG neurons. **a.** Overview of the Cre-mediated gene rearrangements of Rosa26 loci and the transgene *CAG-loxP-stop-LoxP-hNR2F1*. Experimental design is the same used in Figure 2A’. **b.** Histogram showing the linear density of dendritic mitochondria obtained by normalizing the total number of dendritic mitochondria to the total dendritic arborization length in both Ctrl and Nr2f1 -O/E newborn neurons. Unpaired Student’s t test, p=0.4434. **d.** Cumulative frequency distribution for the length of dendritic mitochondria in control vs. Nr2f1 overexpressing mitoDsRed+DCX+YFP+ adult-born neurons. Kolmogoror-Smirnov test, p<0.0002. **d.** Graph reporting the mitochondrial occupancy expressed as the percentage of dendritic length covered by the mitoDsRed signal. Mann Whitney test, p=0.0301. Data are shown as mean±SD. Each dot represents a cell. Data are shown as mean±SD. *p<0.05.

